# Dissociable structural and molecular pathways of age-related variability in sustained attention

**DOI:** 10.64898/2026.06.29.735429

**Authors:** S. Subbulakshmi, Jennifer Park, Julia Rathmann-Bloch, Tyler Ward, Gloria L. Cheng, Douglas S. Miller, Shawn T Schwartz, Jintao Sheng, Tammy T. Tran, Sharon J. Sha, Gayle Deutsch, Alexandra N. Trelle, Elizabeth C. Mormino, Anthony D. Wagner

**Author notes:** **For Correspondence:** S. Subbulakshmi; Anthony D. Wagner.

## Abstract

Sustained attention, the capacity to maintain goal-directed attention over extended periods, declines with age but with substantial individual variability across cognitively unimpaired (CU) older adults. We sought to determine the contributions of microstructural deterioration of the superior longitudinal fasciculus (SLF), the principal white-matter tract coupling prefrontal and parietal nodes of the dorsal attention network, as well as markers of preclinical Alzheimer’s disease (AD) to individual differences in sustained attention in ageing.

In 162 CU older adults (mean (SD) age = 73.0 (4.9), %F = 64.8) drawn from two Stanford ageing cohorts, we measured sustained attention using the gradual-onset Continuous Performance Task (gradCPT). Performance was indexed by a composite score (Att-Z) derived from discriminability (d′) and response-time variability (RTV). We used diffusion MRI tractography to quantify SLF fractional anisotropy (FA) and mean diffusivity (MD) alongside two control tracts, the corticospinal tract (CST) and cingulum (CGC). NULISA multiplex plasma immunoassays were used to measured AD-related pathology (pTau-181, pTau-217), neuroaxonal injury (NfL), and astrocytic reactivity (GFAP; plasma subsample N = 146). Parallel mediation and commonality analyses tested whether structural and plasma biomarker pathways contribute independently to individual differences in sustained attention function.

Older age was associated with lower Att-Z (β = −0.315, p < 0.001). In simultaneous three-tract regression models, only SLF microstructure uniquely predicted Att-Z (FA: β = +0.275, p < 0.001; MD: β = −0.320, p < 0.001). SLF microstructure mediated the age-attention relationship (FA indirect β_unstandardised_ = –0.01, p_boot_ <0.01, MD indirect β_unstandardised_ = –0.009 to –0.01, p_boot_ <0.05). Plasma pTau-181 (β = −0.212, p_FDR_ < 0.03), pTau-217 (β = −0.163, p_FDR_ < 0.05), and NfL (β = −0.156, p_FDR_ < 0.05) were negatively associated with Att-Z and each mediated effects of age on performance (pTau-181 indirect β_unstandardised_ = −0.008, p_boot_ < 0.01; pTau-217 indirect β_unstandardised_= −0.006, p_boot_< 0.05; NfL indirect β_unstandardised_ = −0.011, p_boot_< 0.05), whereas GFAP showed no association with sustained attention in any model. SLF microstructure was negatively associated with age (FA β = −0.207, p < 0.01; MD β = +0.270, p < 0.001) but was not significantly associated with plasma biomarkers (all *p* > 0.10). In parallel mediation models, SLF microstructure and plasma pTau exhibited significant independent indirect effects on Att-Z (SLF FA indirect β_unstandardised_= −0.010, p_boot_< 0.01; SLF MD indirect β_unstandardised_ = −0.009 to −0.010, p_boot_< 0.05; pTau-181 indirect β_unstandardised_ = −0.008 to −0.009, p_boot_< 0.01; pTau-217 indirect β_unstandardised_= −0.006 to −0.007, p_boot_< 0.05). Effects of NfL were attenuated when combined in a model with pTau (indirect β_unstandardised_ = −0.005 to −0.008, p_boot_> 0.15), suggesting shared variance among these markers.

These findings reveal multiple pathways that contribute to individual differences in sustained attention in ageing that are detectable before clinical impairment, including SLF white matter integrity –– which is linked to the structural disconnection of the dorsal attention network –– and preclinical AD pathology. The independence of these pathways identifies two separable biological targets for preserving attentional capacity in CU older adults.

## Introduction

Maintaining goal-directed attention over time is fundamental to everyday cognitive function, yet this capacity declines measurably with age^[1–3]^ and its deterioration forecasts broader functional dependence in later life^[4–6]^. The degree of sustained attention change varies considerably across individuals^[7–11]^, and the neurobiological processes that drive this variability remain poorly understood.

Ageing may influence attentional decline through multiple pathways. First, structural deterioration of the dorsal attention network (DAN) may degrade the neural scaffolding of sustained attention. DAN depends critically on long-range axonal connectivity, with the superior longitudinal fasciculus (SLF) being the principal white matter backbone connecting frontal and parietal nodes of this network^[12,13]^. Extant data indicate that the SLF undergoes progressive microstructural decline later in the adult lifespan^[14–16]^. Second, Alzheimer’s disease (AD)-related amyloid and tau pathology accumulates years to decades before symptom onset in cognitively unimpaired (CU) adults^[17–19]^ and may affect attentional function through multiple pathways, including early involvement of brainstem neuromodulatory systems^[20–22]^ and the involvement of amyloid plaque pathology within frontal and parietal association cortices that support top-down control^[23–25]^. Interestingly, decline in global measures of attention assessed with standard neuropsychological measures often report that Amyloid+ CU show greater decline in this domain compared to Amyloid-CU^[26,27]^. Although these traditional neuropsychological tasks provide insight towards attention in ageing, they are confounded by other processes such as working memory, processing speed, etc. Whether white matter degeneration and preclinical AD pathology represent independent pathways of age-related attentional decline or whether they converge on a common mechanism remains unknown.

The SLF connects lateral prefrontal cortex, including dorsolateral prefrontal and frontal eye field regions, with posterior parietal cortex (superior parietal lobule, intraparietal sulcus, inferior parietal lobule), forming the principal white matter substrate of the DAN that supports top-down, goal-directed visual attention and sustained attentional performance^[12,13,25,28–30]^. Convergent evidence implicates the SLF in attentional function: lesion studies link SLF damage to frontoparietal disconnection and attentional control deficits^[31,32]^, and individual differences in SLF microstructural integrity predict attentional performance across the lifespan^[29,30,32–36]^. Among major white matter pathways, the SLF undergoes particularly pronounced age-related microstructural decline^[14–16]^, and reduced SLF integrity has been associated with poorer attentional ability in older adults^[34–38]^. However, it remains unclear whether SLF microstructure specifically predicts individual differences in sustained attention across CU older adults as well as whether such effects are specific to SLF or reflective of more generalized white matter differences. Demonstrating tract-specific associations would implicate targeted disruption of DAN circuitry, rather than diffuse white matter ageing, as a structural mechanism underlying sustained attentional differences.

Phosphorylated pTau isoforms (pTau-181^[39,42]^ and pTau-217^[40–42]^) are robust biomarkers of underlying AD pathology, with elevations detectable in CU older adults’ years to decades before clinical symptom onset^[17,18,43–45]^. Consistent with work using amyloid PET to capture early AD processes in CU^[27]^, elevated plasma pTau is associated with poorer global cognition^[46,47]^, abstract reasoning^[46]^, and episodic memory^[48]^ in CU older adults, suggesting that preclinical AD pathology detected with plasma biomarkers is associated with effects on cognitive function within the normal range. Although preclinical AD research has disproportionately emphasised memory, converging evidence indicates that early AD pathology is associated with subtle decline across multiple cognitive domains in CU adults, including attention^[26,27]^. Indeed, tau pathology may affect attentional function through disruption of neuromodulatory systems^[20–22,49–52]^ and amyloid pathology is found in frontal and parietal cortices that support top-down attentional control early in CU ^[23–25]^. These associations, however, are typically derived from global attention composites embedded in standard neuropsychological batteries, which conflate several attentional and executive subprocesses rather than isolating a single, well-defined construct. It therefore remains unclear whether plasma markers of AD pathology predict performance on a task that specifically isolates sustained attention in CU older adults, and, if so, whether early AD pathology acts on sustained attention via frontoparietal white matter integrity or is an independent driver of differences in sustained attention. Moreover, it also is unknown whether SLF microstructure and AD pathology capture variance in sustained attention that is distinct from or shared with other markers that are not specific to AD, but change with ageing, such as markers of neuroaxonal damage (e.g., neurofilament light chain [NfL])^[53]^ and/or astrocytic reactivity (glial fibrillary acidic protein [GFAP])^[54]^. Resolving these questions is necessary to characterise the factors that contribute to individual difference in sustained attention in CU older adults.

In the present study, we examined structural and plasma proteomic correlates of sustained attention in CU older adults (61-89 yrs; **Table 1**) drawn from two Stanford cohorts of ageing (DTI sample N = 162; plasma subsample N = 146). Sustained attention was measured using a composite index derived from discriminability and response time variability on the gradual-onset Continuous Performance Task (gradCPT)^[55,56]^, capturing both response accuracy and response consistency effects, respectively, that depend on sustained attentional engagement^[7]^. To index structural integrity, we focused on the SLF within a multi-tract model that permitted assessment of the tract specificity of links to attention. To index AD pathology and additional non-specific markers, we examined plasma pTau-181 and pTau-217 alongside NfL, a marker of non-specific neuroaxonal damage that rises with age across multiple etiologies^[53,57,59]^, and GFAP, a marker of astrocytic reactivity^[54,58,60]^. Our first objective was to determine whether SLF microstructural integrity specifically predicts individual differences in sustained attention and mediates age-related attentional variability, with effects exceeding those of control white matter tracts. Our second objective was to determine whether plasma pTau-181, pTau-217, NfL, and/or GFAP predict individual differences in sustained attention and mediate age-related attentional variability, and whether such effects are independent or related. Our third objective was to determine whether SLF microstructure and plasma biomarkers account for independent variance in sustained attention or whether they converge on a shared pathway linking ageing to attentional decline through preclinical AD.

**Table 1:**
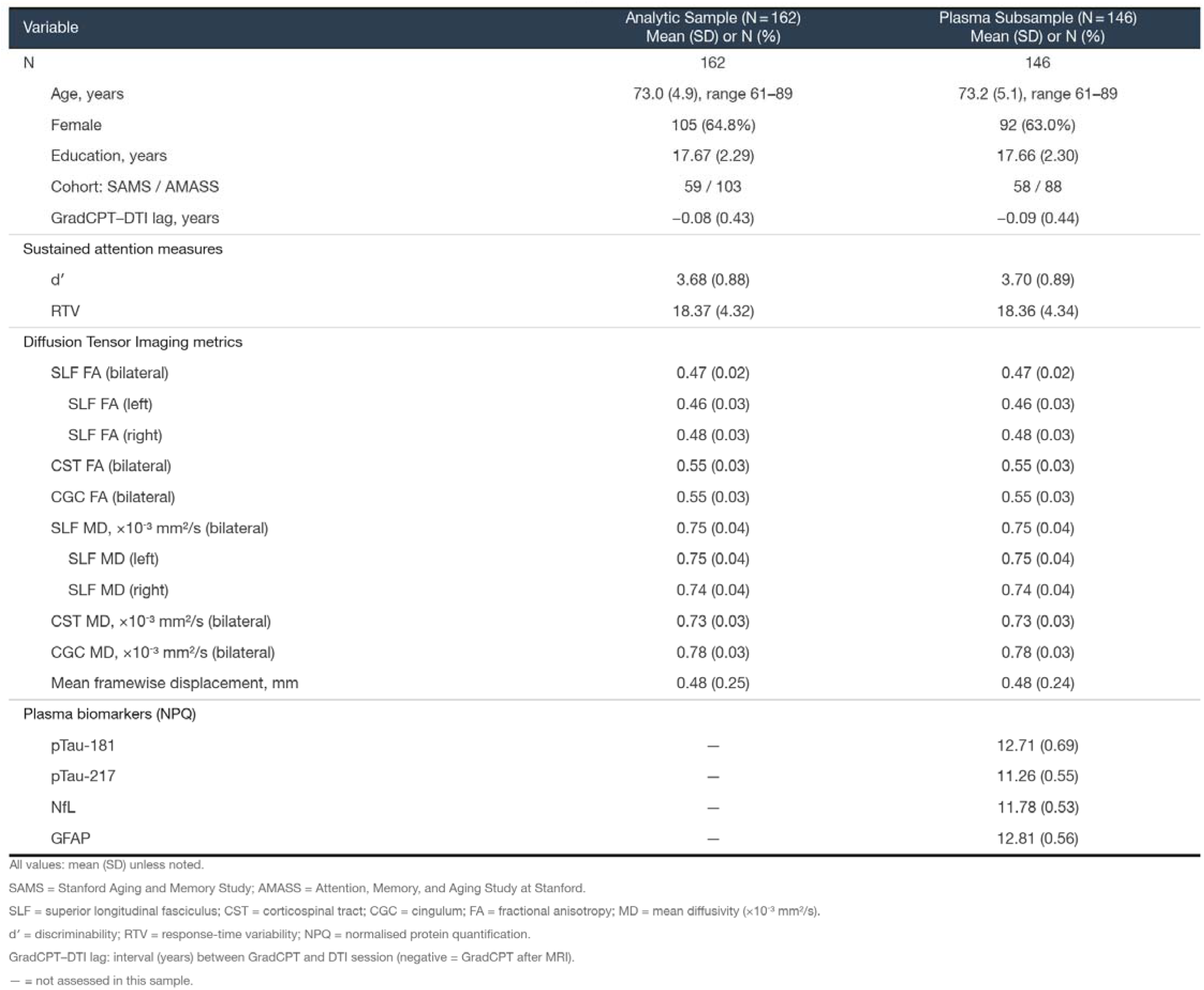
Demographics and descriptive statistics for the full DTI analytic sample (N=162). Age, education, and continuous measures are presented as mean (SD); sex and cohort as N (%). Sustained attention performance was assessed using the gradual-onset Continuous Performance Task (gradCPT): discriminability (d′) indexes the ability to distinguish targets from non-targets; response-time variability (RTV) reflects within-person across-trial variability in reaction time. gradCPT–DTI lag reflects the interval between cognitive testing and DTI acquisition (negative = DTI preceded testing). DTI metrics are bilateral tract-averaged or hemisphere-specific FA and MD values derived from PyAFQ tractography following QSIPrep preprocessing. Mean framewise displacement indexes in-scanner head motion. Plasma biomarkers of interest (pTau-181, pTau-217, NfL, GFAP) were measured using NULISAseq assays and are reported in NPQ units.

Foreshadowing the results, we demonstrate that SLF microstructural integrity specifically predicts sustained attention, with effects exceeding those of control tracts, and mediates effects of age on sustained attention. Plasma pTau-181 and pTau-217 each predicts sustained attention and mediates age-related variability, effects that remain significant when examined jointly with SLF microstructure. While NfL also is linked to attentional variability, independent of SFL, this factor was no longer significant when modelled alongside pTau-181, whereas GFAP shows no association with attention. Collectively, these findings reveal dissociable pathways that explain heterogeneity in sustained attentional capacity in CU adults, and lay the groundwork for identifying individuals at risk of accelerated attentional decline in the preclinical phase of AD.

## Methods

### Participants

Participants were 162 cognitively unimpaired (CU) older adults (mean age = 73.0 ± 4.9 yrs; 64.8% female; education = 17.67 ± 2.29 yrs; **Table 1**) drawn from the Stanford Aging and Memory Study (SAMS; N = 59) and the Attention, Memory, and Aging Study at Stanford (AMASS; N = 103). In SAMS, cognitively unimpaired status was determined via a Clinical Dementia Rating (CDR) score of 0, performance within 1.5 standard deviations of demographically adjusted means on standardised neuropsychological assessments, and a consensus diagnosis of CU by a panel of neurologists and neuropsychologists ^[61,62]^. In AMASS, cognitively unimpaired status was determined based on (a) a score greater than 26 (out of 30) on the telehealth version of the MoCA and (b) performance within 1.5 standard deviations of age-adjusted norms on an in-person neuropsychological battery similar to that administered in SAMS. Additional eligibility criteria for both studies included normal or corrected-to-normal vision and hearing, right-handedness, native English speaking, and no history of neurological or psychiatric disease. All participants provided written informed consent and were compensated for their time, following procedures approved by the Stanford University Institutional Review Board.

All participants (N = 162) had diffusion tensor imaging (DTI) MRI and sustained attention data and formed the primary analytic sample for white-matter analyses; N = 161 had usable data for all three DTI tracts of interest (see below) and were included in the three-tract specificity model. Moreover, a subset (N = 146) had plasma biomarker data and were included in plasma analyses and in joint mediation/commonality models combining DTI and plasma predictors.

### Sustained-Attention Task

Sustained attention was assessed using the gradual-onset Continuous Performance Task (gradCPT^[7,55,56]^). Across the task, city and mountain scene images transitioned gradually (each image fading in over 800 ms while the previous image faded out), and participants responded with a button press to frequent city scenes (90% of trials) and withheld responses to rare mountain scenes (10% of trials). The task yielded two primary participant-level indices of sustained attention: (a) discriminability (d’), calculated using signal-detection theory as z(hit rate) − z(false-alarm rate), indexing the ability to distinguish go-stimuli from no-go-stimuli; and (b) mean response-time variability (RTV), calculated as the mean of the coefficient of variation (standard deviation divided by the mean) of reaction times on correct go-trials, indexing moment-to-moment fluctuations in attentional engagement^[46]^. To confirm that findings were not specific to either measure in isolation, we also computed an attention composite score, Att-Z = (z_d′ − z_RTV) / 2, where higher values indicate better overall attentional performance. We report all three metrics for the regression models, and the mediation and variance-partitioning analyses focus on Att-Z.

### DTI MRI Acquisition, Preprocessing, and Tractography

Diffusion-weighted MRI was acquired on a GE PET-MR scanner (3T; 80 slices were acquired with 2×2×2 mm^3^ resolution and 30 diffusion directions [3 b0s, b-value=1000 s/mm2]). A ‘flip’ sequence was collected prior to diffusion imaging to enable field map correction.

Diffusion images were preprocessed with QSIPrep^[63]^, which applies susceptibility distortion correction, eddy-current correction, motion correction, and b0-field normalization within a unified workflow. Head motion was quantified as mean framewise displacement (FD; mm) and included as a covariate in all DTI-based regression models.

Tract segmentation and quantification were performed using pyAFQ^[64,65]^, an automated white-matter tract identification pipeline that fits deterministic streamline tractography and segments tracts using atlas-based waypoint regions of interest. Of the full pyAFQ tract set, three tracts were of primary interest: the superior longitudinal fasciculus (SLF), the principal frontoparietal association bundle connecting the cortical nodes of the dorsal attention network^[13]^; the corticospinal tract (CST), a predominantly motor pathway that was included as an anatomically distinct comparison tract^[66]^; and the cingulum (CGC), a limbic association bundle^[67]^ not typically implicated in goal-directed sustained attention that was included as an additional anatomically distinct comparison tract. For each tract (SLF, CST, CGC; for tractography depictions see **Supplementary Section S1**), left and right hemisphere tracts were averaged into a bilateral mean as the primary measure (hemispheric analyses are reported in **Supplementary Section S1**). For each tract, fractional anisotropy (FA) and mean diffusivity (MD) were computed as the median value across streamline nodes. Data from one subject did not yield converging CST FA values and were excluded from corresponding FA and MD analyses that included CST.

### Plasma Biomarkers

Plasma samples from 146 participants (**Table 1**) were included in the current study (58 from SAMS and 88 from AMASS). Plasma collection and storage methods were harmonized across cohorts. EDTA plasma was collected by venipuncture, centrifuged for 10 min at 2000 x g, aliquoted in polypropylene tubes, and stored at −80°C until measurement. Plasma samples were analysed using the NULISAseq CNS panel as described previously^[51]^. Here we include four a priori plasma targets of interest: pTau-181 and pTau-217 (AD-specific phosphorylated tau isoforms), GFAP (glial fibrillary acidic protein; astrocytic injury / reactive astrogliosis), and NfL (neurofilament light chain; neuroaxonal injury). All plasma concentrations are reported in NULISA Protein Quantification (NPQ) units. NPQ units reflect log2-transformed counts normalized to within-well internal controls and plate-specific target medians.

### Statistical Analysis

All analyses were conducted in Python (v 3.9) using statsmodels and SciPy. Ordinary least-squares (OLS) regression was used throughout, with standardised beta coefficients (β) reported alongside unstandardised coefficients (b) where informative. Given strong a priori predictions about the direction of hypothesized relationships, one-tailed thresholds were adopted accordingly.

#### Regression covariates

All linear regressions controlled for age (whenever age was not a predictor of interest), sex, and education. DTI models additionally controlled for mean framewise displacement (FD). Mediation models and commonality analyses used the same covariate set.

#### FDR correction

For families of plasma biomarker tests (age → biomarker; biomarker → attention), p-values were corrected for multiple comparisons using the Benjamini–Hochberg procedure^[68]^, applied separately within each outcome family. We report non-corrected p-values as p and FDR corrected values as p_FDR_.

#### Tract specificity

To isolate SLF-specific explained variance in attention performance, FA or MD values from the SLF, CST, and CGC were entered simultaneously into the same regression model, so that each tract’s coefficient reflects unique variance after partialling out the others. Variance inflation factors were inspected to confirm acceptable multicollinearity (all VIF < 1.5). Bootstrap comparisons (5,000 resamples) tested whether the SLF FA or MD standardised beta differed significantly from those of the comparison tracts; one-tailed p-values were derived from the bootstrap distribution of Δβ.

#### Mediation analyses

Mediation analyses used bootstrap resampling (5,000 samples) to estimate indirect effects (a × b path product) and bias-corrected 90% confidence intervals. The a-path modelled the predictor-to-mediator association; the b-path modelled the mediator-to-outcome association controlling for the predictor and covariates. Significance for the indirect effect was assessed by whether the 90% CI excluded zero and by the one-tailed bootstrap p-value. The CI was 90% to be consistent with the one-tailed boot-strapped distributions. Parallel mediation models (Hayes Model 4^[69]^) entered two mediators (i.e., one structural and one molecular) simultaneously to test whether each explained unique indirect variance after partialling out the other. Boot-strapped p-values are presented as p_boot_.

#### Commonality analysis

Based on observed relationships between plasma biomarkers and attention (see Results below), to formally decompose the shared and unique contributions of age, SLF FA or MD, and plasma biomarkers to attentional outcomes, we leveraged commonality analysis^[70,71]^, which uses the inclusion– exclusion principle to partition the total R^2^ of a predictor set (above the other covariates) into unique and shared variance components. For three predictors, this yields seven non-overlapping components (three unique, three pairwise shared, one triple-shared); negative shared values indicate suppression. Unique variance components were tested using one-tailed OLS p-values from the joint regression model including all three predictors simultaneously. Shared variance components, which have no tractable parametric null distribution, were assessed using one-tailed bootstrap p-values (5,000 resamples), with significance defined as p < .05.

## Results

### 1. Sustained Attention Performance

#### 1.1 Age and sustained attention performance relationships

Older age was associated with lower gradCPT discriminability (d′; β = −0.314, p < 0.001; **Fig. 1A**), greater response-time variability (RTV; β = +0.274, p < 0.001; **Fig. 1B**), and lower attention composite score (Att-Z; β = −0.315, p < 0.001; **Fig. 1C**; where higher values of Att-Z reflect better sustained attention performance). These results confirm robust age-related decline in sustained attention across both constituent measures and the composite, providing the primary effect to be mechanistically decomposed in subsequent analyses.

**Figure 1.**
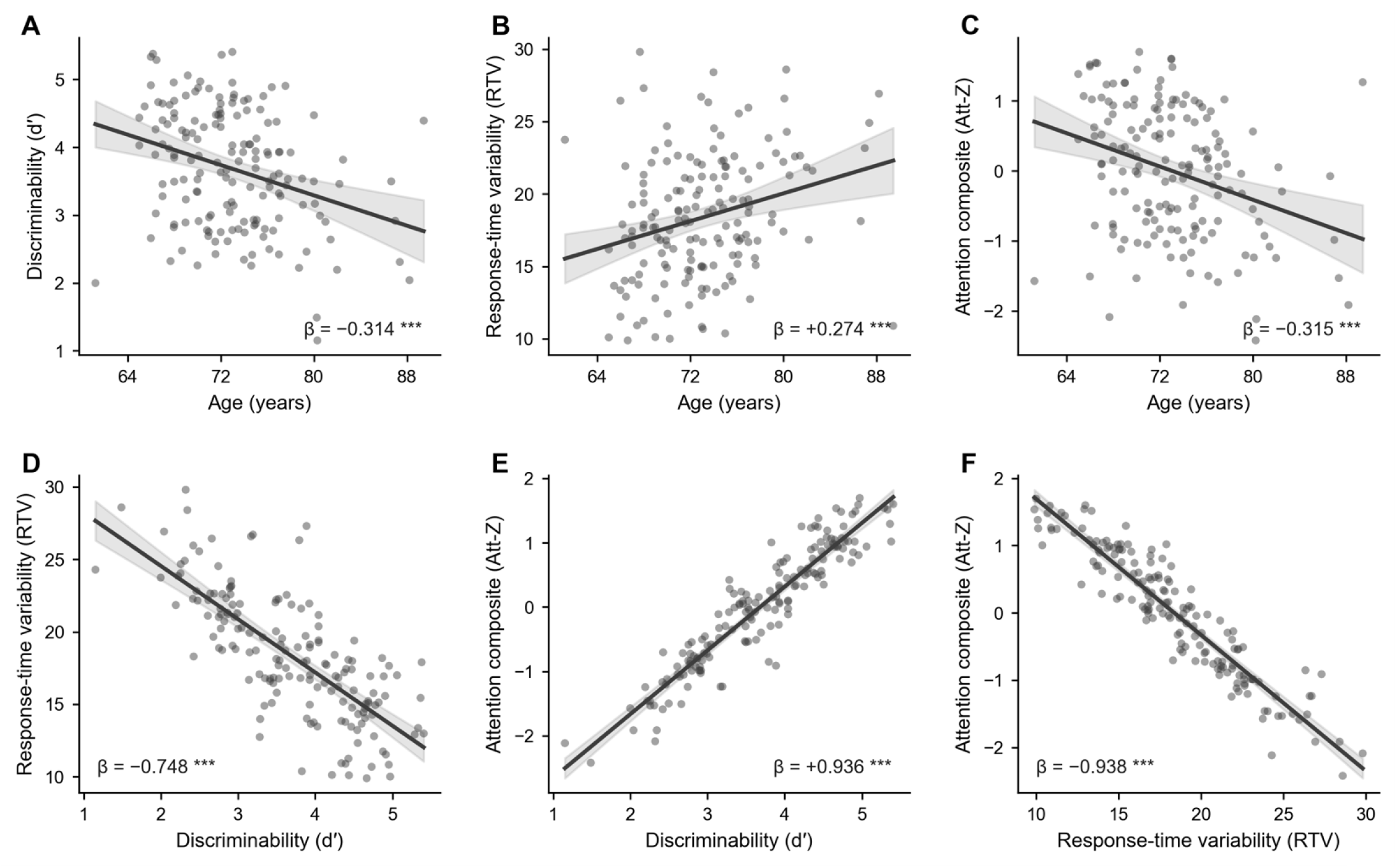
Age and sustained attention performance relationships (N = 162). Top row: Age associations with gradCPT discriminability (A), gradCPT response-time variability (B), and the attention composite score (C). Bottom row: Pairwise correlations among attention measures (D–F). Lines show model-predicted regression fits with 95% CI; β annotations control for sex and education. All p-values are one-tailed. ***p < 0.001.

The three attention measures were highly intercorrelated (d′–RTV: partial r = −0.748; d′–Att-Z: r = +0.936; RTV–Att-Z: r = −0.938; all p < 0.001; **Fig. 1D–F**), confirming that Att-Z captures shared variance across both constituent measures (as would be expected given that it is a composite of d′ and RTV).

### 2. White-Matter Microstructure

#### 2.1 SLF Microstructure Is Associated with Age

Older age was associated with lower bilateral superior longitudinal fasciculus (SLF) FA (β = −0.207, p < 0.01; **Fig. 2A**) and higher SLF MD (β = +0.270, p < 0.001; **Fig. 2E**), confirming age-related decline in SLF microstructural integrity. The pattern of age-related FA decline differed numerically across tracts: while significant for SLF, it was marginal for CGC (β = −0.127, p = 0.052), and absent for CST (β = +0.009, p = 0.917). However, the overall age × tract interaction did not reach significance (F(2,504) = 1.73, p = .178), with only a marginal SLF–CST difference (p = .083). By contrast, age-related MD increases were consistent across all three tracts (for CST: β = +0.335 and CGC: β = +0.271, both p < .001), and there was no age × tract interaction (F(2,504) = 0.022, p = 0.978).

**Figure 2.**
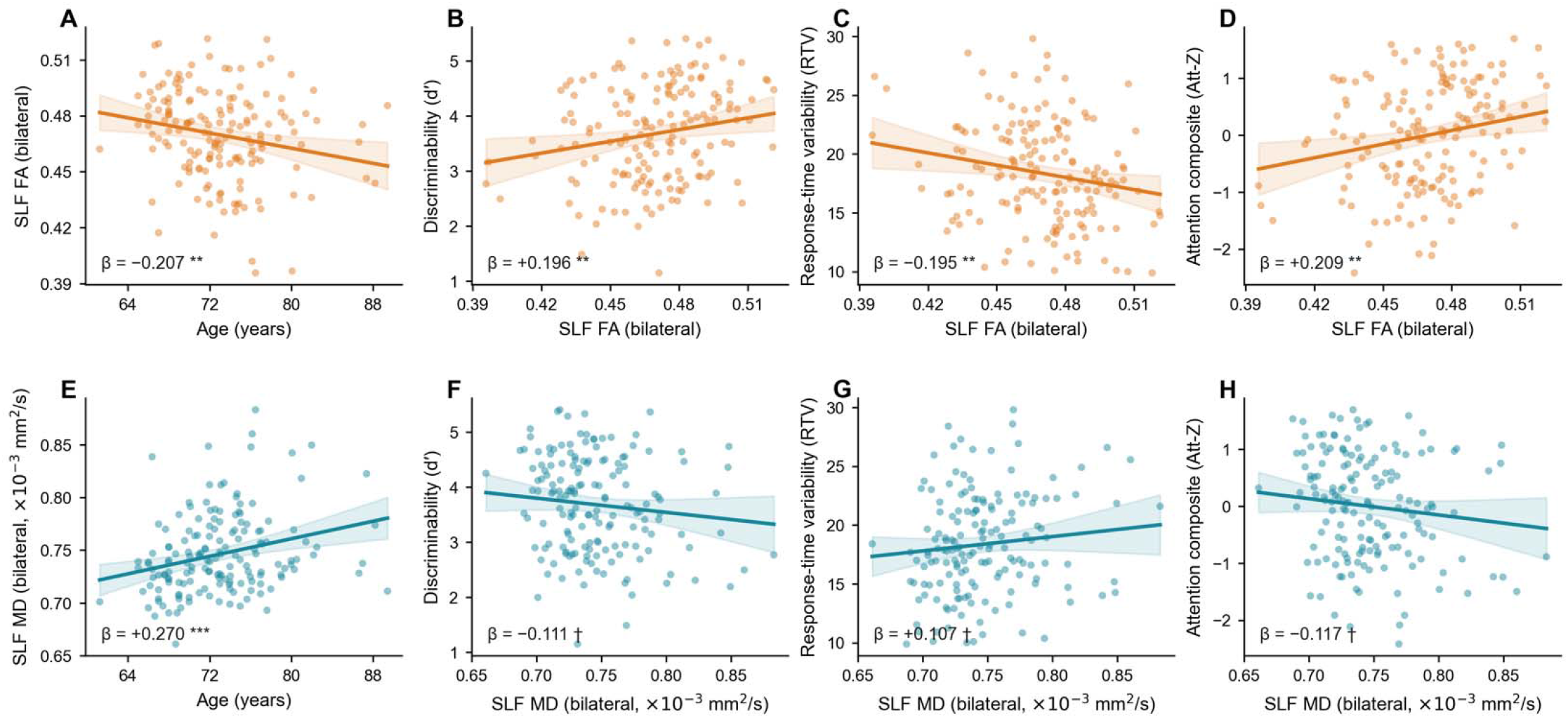
Partial-regression associations (N=162) between bilateral SLF fractional anisotropy (FA; orange, top row) or mean diffusivity (MD; teal, bottom row) and age (A, E) or sustained attention (B-D, F-H). Older age was associated with lower SLF FA and higher SLF MD. Higher SLF FA and lower MD were associated with better gradCPT discriminability (d′; B, F), lower response-time variability (RTV; C, G), and a higher attention composite score (Att-Z; D, H). Each panel plots observed outcome values against the Age or DTI predictor after residualising both variables on covariates. Lines and shading show OLS predicted values ± 95% CI. Standardised β coefficients and significance symbols refer to the partial regression of the Age or DTI metric on the outcome. †p < 0.10, **p < 0.01, ***p < 0.001 (one-tailed).

#### 2.2 SLF Microstructure Predicts Sustained Attention

We next tested whether SLF microstructural integrity predicted sustained attention, and whether any such association was specific to the SLF relative to the CST and CGC. Higher bilateral SLF FA was associated with better gradCPT performance: SLF FA positively predicted d′ (β = +0.196, p < 0.01; **Fig. 2B**), negatively predicted RTV (β = −0.195, p < 0.01; **Fig. 2C**), and positively predicted the attention composite (Att-Z; β = +0.209, p < 0.01; **Fig. 2D**). Consistent with these FA findings, lower SLF MD showed directionally consistent but marginally significant associations with attention: SLF MD negatively predicted d′ (β = −0.111, p = 0.081; **Fig. 2F**), positively predicted RTV (β = +0.107, p = 0.094; **Fig. 2G**), and negatively predicted Att-Z (β = −0.117, p = 0.072; **Fig. 2H**). For analyses of hemispheric SLF microstructure relationships with attention, see **Supplementary Section S1**.

To test whether sustained attention is differentially related to SLF integrity (relative to the comparison tracts), FA from the SLF, CST, and CGC were entered simultaneously into the same regression (N = 161; one subject without converging CST FA values was excluded). Importantly, SLF FA uniquely predicted all three attention measures: d′ (β = +0.273, p < 0.01; **Fig. 3A**), RTV (β = −0.240, p < 0.01; **Fig. 3B**), and Att-Z (β = +0.275, p < 0.01; **Fig. 3C**). By contrast, CST showed no significant unique associations with any attention outcome (all p > 0.09) and CGC FA reached significance for RTV (β = +0.168, p < 0.05; **Fig 3B**) and Att-Z (β = −0.138, p < 0.05; **Fig 3C**), but in the direction inconsistent with a white-matter integrity interpretation, suggesting suppression by the correlated tracts (VIF < 1.5 for all predictors). Bootstrap comparisons confirmed that the SLF FA coefficient was significantly stronger than that for CST and CGC for d′ (SLF vs. CST: Δβ = +0.376, p_boot_ < 0.005; vs. CGC: Δβ = +0.362, p_boot_ < 0.005), RTV (vs. CST: Δβ = −0.287, p_boot_ < 0.05; vs. CGC: Δβ = −0.411, p_boot_ < 0.01), and Att-Z (vs CST: Δβ = +0.355, p_boot_ < 0.005; vs CGC: Δβ = +0.414, p_boot_ < 0.005).

**Figure 3.**
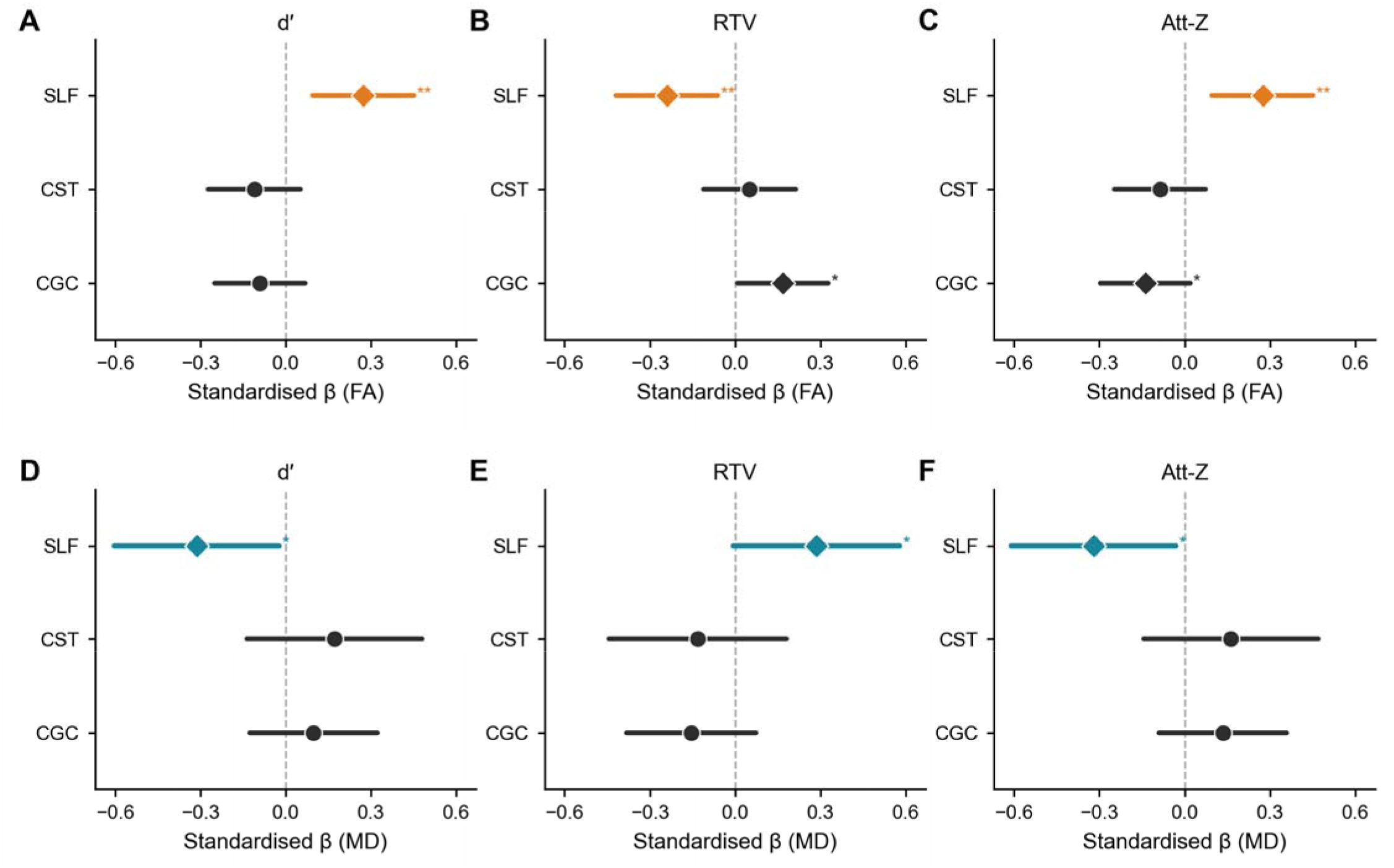
Tract specificity of white-matter associations with sustained attention, revealing unique variance explained by SLF (N = 161). Coefficient plots show standardised β estimates and 95% CIs from simultaneous OLS models in which FA (top row, A–C) or MD (bottom row, D–F) from the superior longitudinal fasciculus (SLF), corticospinal tract (CST), and cingulum (CGC) were entered as simultaneous predictors of d′ (A, D), RTV (B, E), and Att-Z (C, F), controlling for age, sex, education, and mean framewise displacement. Each tract’s coefficient reflects explained variance unique to that tract after partialling out the other two. Diamond markers indicate p < 0.05; circle markers indicate n.s. *p < 0.05, **p < 0.01 (one-tailed). SLF coefficients were significantly larger than those of CST and CGC across all outcomes (bootstrap comparisons in main text).

A parallel pattern was observed using MD (**Fig. 3D–F**): SLF MD uniquely predicted d′ (β = −0.312, p < 0.05), RTV (β = +0.286, p < 0.05), and Att-Z (β = −0.320, p < 0.05); CST and CGC MD were non-significant (all p > 0.09). Bootstrap comparisons confirmed that SLF MD associations were significantly stronger than that for CST and CGC for d′ (vs. CST: Δβ = −0.477, p_boot_ < 0.01; vs. CGC: Δβ = −0.407, p_boot_ < 0.01) and Att-Z (vs. CST: Δβ = −0.473, p_boot_ < 0.05; vs. CGC: Δβ = −0.448, p_boot_ < 0.05). For RTV, SLF MD was significantly stronger than CGC MD (Δβ = +0.431, p_boot_ < 0.05), with the CST comparison trending in the expected direction (Δβ = +0.410, p_boot_ = 0.062). These analyses indicate that SLF microstructure also explains unique variance in sustained attention across CU older adults (controlling for age, sex, and education) and that this association is specific to the SLF rather than reflecting diffuse white-matter microstructural differences.

### 3. Plasma Biomarkers

#### 3.1 Age Associations and Intercorrelations of the Plasma Biomarkers

Age was associated with higher concentrations of pTau-181 (β = +0.202, p < 0.01, p_FDR_ < 0.01), pTau-217 (β = +0.212, p < 0.01, p_FDR_ < 0.01), NfL (β = +0.386, p < 0.001, p_FDR_ < 0.001), and GFAP (β = +0.333, p < 0.001, p_FDR_ < 0.001) (**Fig 4**). Correlations among the four plasma biomarkers were low to strong, with the two pTau markers being most tightly coupled (r = 0.673) and GFAP and pTau-181 being most distinct (r = 0.181) (**Fig 4E**).

**Figure 4.**
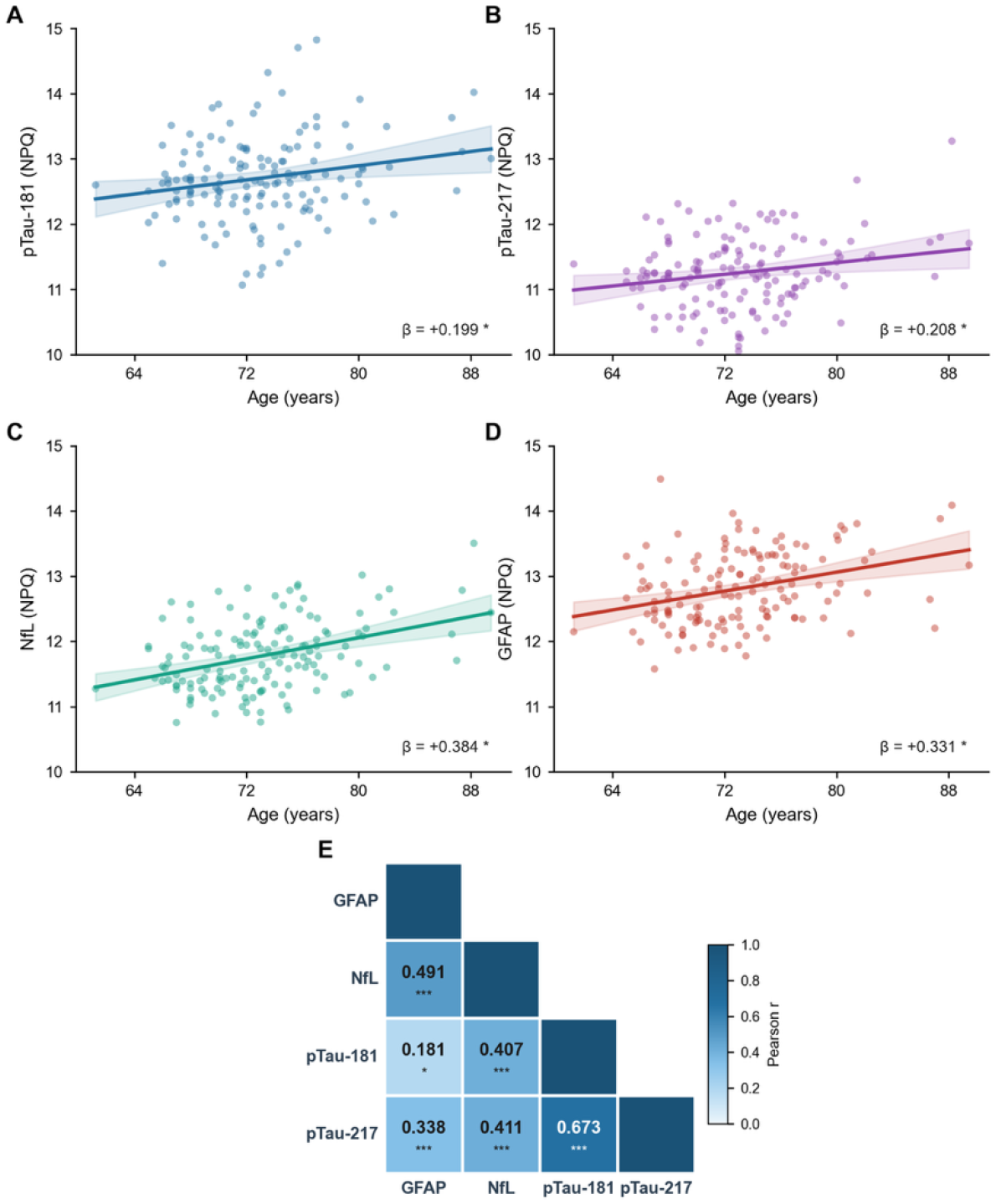
Age associations with primary plasma biomarkers and plasma intercorrelations (N = 146). Scatter plots show age vs. pTau-181 (A), pTau-217 (B), NfL (C), and GFAP (D) with model-predicted regression line and 95% CI. β values control for sex and education. p-values are one-tailed; *FDR-corrected p_FDR_ < 0.05. (E) Pairwise correlations (Pearson r values) among the four plasma biomarkers, *p < 0.05, ***p < 0.001 (one-tailed).

#### 3.2 Associations Between Plasma Biomarkers and Sustained Attention

Sustained attention performance declined with increasing pTau-181, pTau-217, and NfL levels. Specifically, gradCPT d′ (**Fig. 5A–D**) was negatively associated with pTau-181 (β = −0.228, p < 0.01, p_FDR_ < 0.02) and NfL (β = −0.184, p < 0.02, p_FDR_ < 0.04); pTau-217 showed a marginal relationship (β = −0.126, p < 0.07, p_FDR_ < 0.09), whereas GFAP was not significantly associated with d′ (p > 0.38). GradCPT RTV (**Fig. 5E–H**) showed nominally significant positive associations with pTau-181 (β = +0.168, p < 0.03, p_FDR_ < 0.06) and pTau-217 (β = +0.178, p < 0.02, p_FDR_ < 0.06); NfL and GFAP were not significantly associated with RTV (both p > 0.11). Finally, the attention composite Att-Z (**Fig. 5I–L**) showed significant negative associations with pTau-181 (β = −0.212, p < 0.01, p_FDR_ < 0.03), pTau-217 (β = −0.163, p < 0.03, p_FDR_ < 0.05), and NfL (β = −0.156, p < 0.04, p_FDR_ < 0.05), whereas GFAP was not associated with Att-Z (p > 0.38).

**Figure 5.**
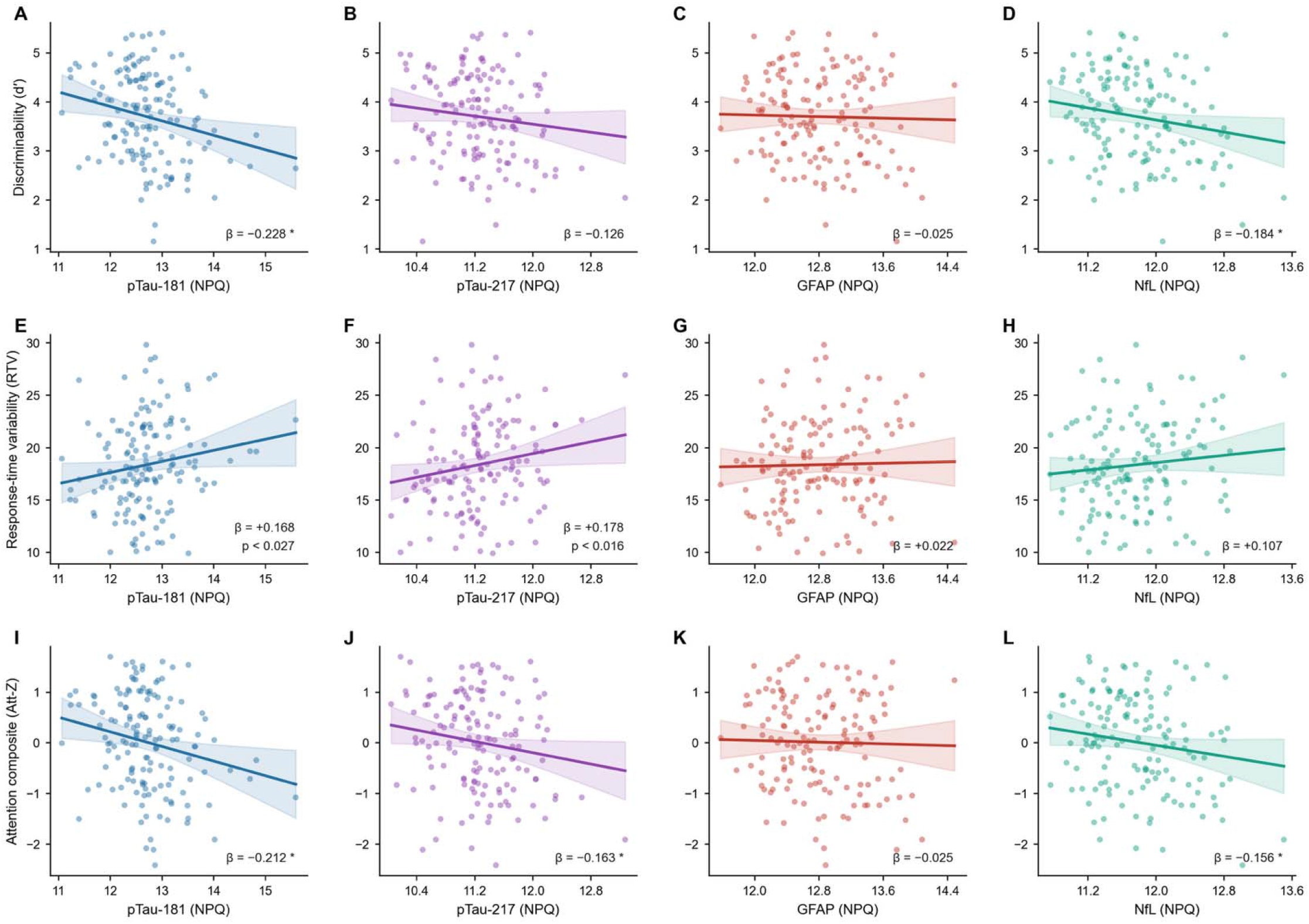
Associations between plasma biomarkers and sustained attention measures (N = 146). Rows show associations with gradCPT discriminability (A–D), response-time variability (E–H), and attention composite score (I–L). Columns show pTau-181, pTau-217, NfL, and GFAP. Lines show model-predicted regression fits with 95% CI; β annotations control for age, sex, and education. All p-values are one-tailed. * p_FDR_ < 0.05 (FDR-corrected); uncorrected p-values shown where p < 0.05.

#### 3.3 Plasma Biomarkers Do Not Predict SLF Microstructure

None of the four plasma biomarkers (pTau-181, pTau-217, NfL, GFAP) significantly predicted bilateral SLF FA (all p > 0.10; **Fig. S3**). Similarly, for SLF MD, GFAP showed a marginal association (β = +0.119, p = 0.078, p_FDR_ >0.05) that did not survive FDR correction and all other biomarkers were non-significant (all p > 0.10). Full statistics and figures are provided in **Supplementary Section S2**.

When combined with the above reported relationships between (a) SLF microstructure and attention and (b) pTau-181, pTau2-17, and NfL burden and attention, these null relationships between plasma biomarkers and SLF microstructural integrity raise the possibility that SLF microstructure and plasma AD pathological burden represent dissociable structural and molecular pathways that drive differences in sustained attention.

### 4. Parallel Mediation: Independent Pathways of Age-Related Attentional Decline

To formally test whether SLF white-matter microstructure and plasma AD pathology represent independent pathways of age-related attentional decline, we estimated parallel mediation models with SLF microstructure and plasma biomarkers (restricted to the three plasma biomarkers that showed FDR-corrected significant associations with Att-Z: pTau-181, pTau-217, and NfL (all p_FDR_ < 0.05; **Section 3**)) as simultaneous mediators of the Age → Att-Z relationship. Each pairwise model included bilateral SLF FA (or SLF MD) and one plasma biomarker as simultaneous mediators, controlling for sex, education, and mean framewise displacement.

#### Pairwise models

SLF FA showed an independent indirect effect on Att-Z in all three pairwise models (pTau-181 model: SLF FA indirect β = −0.010, 90% CI [−0.018, −0.003]; pTau-217 model: SLF FA indirect β = −0.010, 90% CI [−0.018, −0.003]; NfL model: SLF FA indirect β = −0.010, 90% CI [−0.019, −0.003]; all p_boot_ < 0.01, **Fig. 6A–C**). Moreover, all three plasma biomarkers showed a significant independent indirect effect alongside SLF FA (pTau-181: indirect β = −0.0084, 90% CI [−0.016, −0.002], p_boot_ < 0.01; pTau-217: indirect β = −0.0064, 90% CI [−0.015, 0.000], p_boot_ < 0.05; NfL: indirect β = −0.0111, 90% CI [−0.023, 0.002], p_boot_ < 0.05). Parallel models using SLF MD yielded consistent results: SLF MD retained significant indirect effects in all three pairwise models (pTau-181 model: SLF MD indirect β = −0.010, 90% CI [−0.019, −0.002]; pTau-217 model: SLF MD β = −0.009, 90% CI [−0.018, −0.001]; NfL model: SLF MD β = −0.009, 90% CI [−0.018, −0.001]; all p_boot_ < 0.05; **Fig. 6D–F**), and pTau-181 (indirect β = −0.009, 90% CI [−0.017, −0.002], p_boot_< 0.01), pTau-217 (indirect β = −0.007, 90% CI [−0.015, 0.000], p_boot_ < 0.05), and NfL (indirect β = −0.010, 90% CI [−0.022, −0.001], p_boot_ < 0.05) each showed significant independent indirect effects alongside SLF MD.

**Figure 6.**
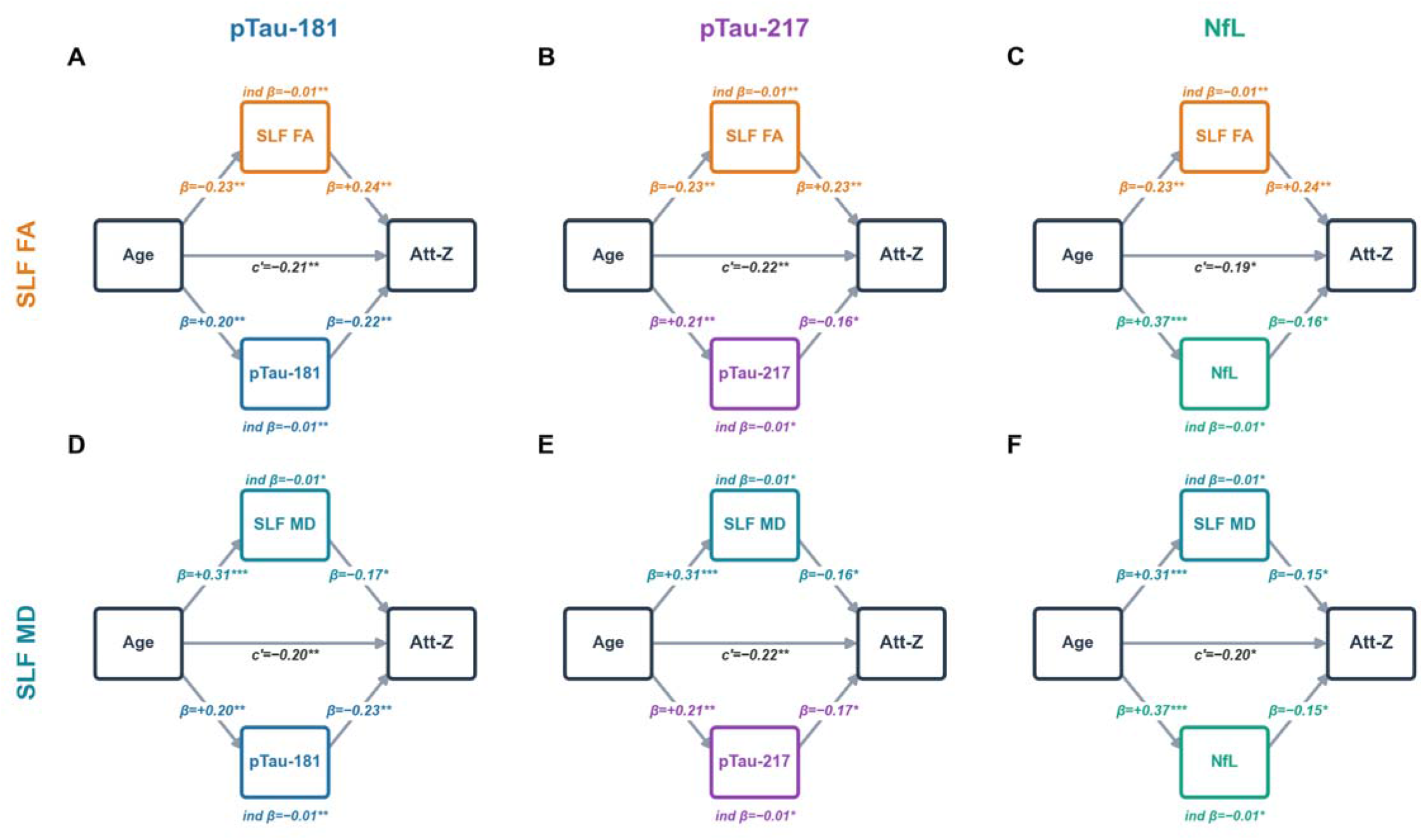
Parallel mediation of age-related attentional decline by SLF white-matter microstructure and plasma biomarkers (N=146). Path diagrams illustrating parallel pairwise mediation models in which age simultaneously acts through bilateral SLF fractional anisotropy (FA; panels A–C) or mean diffusivity (MD; panels D–F) and one plasma biomarker (columns: pTau-181, pTau-217, NfL) to predict the attention composite (Att-Z). Standardised β coefficients are shown on each path; c′ denotes the direct effect of age on Att-Z after accounting for both mediators. Indirect effects (ind β) are unstandardised bootstrap estimates (5,000 resamples). All models controlled for sex, education, and mean framewise displacement. *p or p_boot_ < 0.05, **p or p_boot_ < 0.01, ***p or p_boot_ < 0.001.

#### Full models

To assess the independence of the structural and molecular pathways and the specificity of AD pathological effects relative to neuroaxonal injury, we estimated a full parallel mediation model simultaneously including bilateral SLF FA, pTau-181, and NfL. SLF FA retained a significant independent indirect effect of age on Att-Z (indirect β = −0.0087, 90% CI [−0.017, −0.002], p_boot_ < 0.01), as did pTau-181 (indirect β = −0.0068, 90% CI [−0.015, −0.001], p_boot_ < 0.05), while the NfL effect was attenuated and no longer significant (indirect β = –0.008, 90% CI [−0.021, +0.004], p_boot_ > 0.15); the latter suggests shared variance between NfL and pTau-181, rather than an independent non-AD pathway. An equivalent model using SLF MD yielded consistent results: SLF MD (indirect β = −0.0099, 90% CI [−0.019, −0.002], p_boot_ < 0.05) and pTau-181 (indirect β = −0.0078, 90% CI [−0.015, −0.002], p_boot_ < 0.05) retained significant independent indirect effects, while NfL was again attenuated (indirect β = −0.005, 90% CI [−0.016, +0.005], p_boot_ > 0.23).

Equivalent parallel mediation models with pTau-217 instead of pTau-181 revealed that SLF microstructure again retained significant independent indirect effects (SLF FA: indirect β = −0.010, 90% CI [−0.018, −0.003], p_boot_ < 0.01; SLF MD: indirect β = −0.009, 90% CI [−0.019, −0.001], p_boot_ < 0.05). By contrast, pTau-217 (FA model: indirect β = −0.005, 90% CI [−0.013, +0.001], p_boot_ = 0.076; MD model: indirect β = −0.005, 90% CI [−0.014, +0.001], p_boot_ = 0.066) and NfL (FA model: indirect β = −0.008, 90% CI [−0.019, +0.001], p_boot_ = 0.092; MD model: indirect β = −0.007, 90% CI [−0.021, +0.003], p_boot_ = 0.138) were attenuated to non-significance in both models. While these outcomes suggest that pTau-217 and NfL explain overlapping variance that is more difficult to disentangle in a joint model in this sample, the pTau-181 models provide evidence for separable SLF microstructural and AD-specific molecular pathways of age-related attentional decline, over and above shared variance with NfL.

### 5. Shared and Unique Variance: Commonality Analysis

To complement the above mediation framework, we applied commonality analysis to partition the unique and shared contributions of age, SLF microstructure, and each FDR-significant plasma biomarker to Att-Z across all six DTI × biomarker combinations. Combined models explained 12.8–18.1% of variance in Att-Z (all p < 0.001; **Fig. 7**).

**Figure 7.**
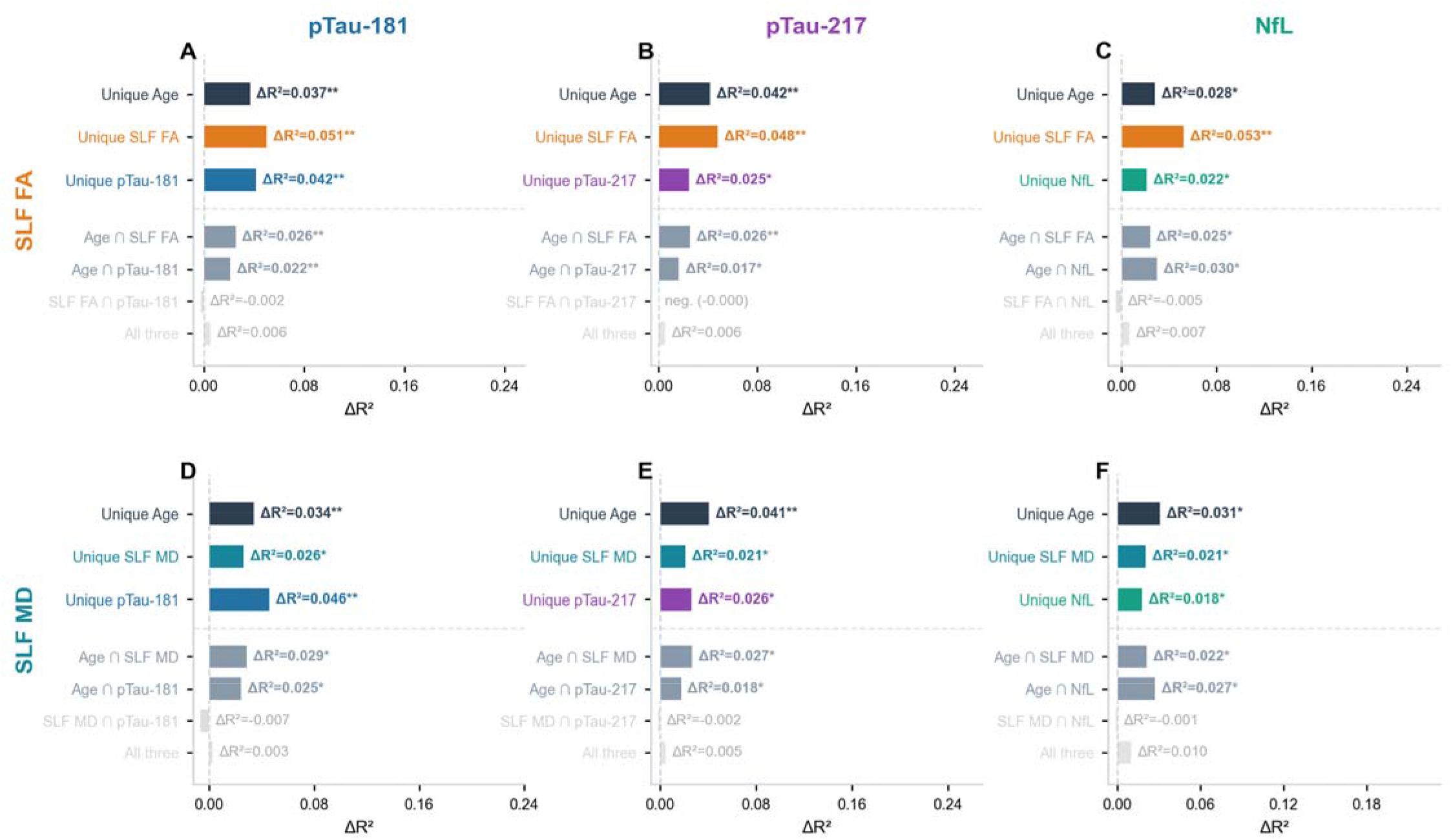
Commonality analysis partitioning explained variance (ΔR^2^) in the composite attention score (Att-Z) among age, SLF white-matter microstructure (FA: top row; MD: bottom row), and each plasma biomarker (pTau-181, pTau-217, NfL; columns A–F), covarying for sex, education, and mean framewise displacement. Bars show unique contributions of each predictor and pairwise/triple shared components. Opaque bars indicate p < .05; faded bars are non-significant. Unique component significance is based on one-tailed OLS p-values (p) from the joint regression model; shared component significance is based on one-tailed bootstrapped p-values (p_boot_) (5,000 resamples). * p or p_boot_ < .05, ** p or p_boot_ < .01.

In pTau-181 models, all three predictors explained significant unique variance (age: ΔR^2^ = 0.034–0.037, p < 0.02; SLF FA/MD: ΔR^2^ = 0.027–0.051, p < 0.03; pTau-181: ΔR^2^ = 0.042–0.046, p < 0.01). Similarly, in pTau-217 models, all three predictors explained significant unique variance (age: ΔR^2^ = 0.041–0.042, p < 0.02; SLF FA/MD: ΔR^2^ = 0.021-0.048, p < 0.05; pTau-217: ΔR^2^ = 0.025–0.026, p < 0.03). In NfL models, age was significant (ΔR^2^ = 0.028–0.031, p < 0.03) and SLF FA was the dominant white-matter predictor (ΔR^2^ = 0.053, p < 0.01); SLF MD also reached significance (ΔR^2^ = 0.021, p < 0.05). In NfL models, all three predictors explained significant unique variance: age (ΔR^2^ = 0.028–0.031, p < 0.02), SLF FA/MD (ΔR^2^ = 0.021-0.053, p < 0.05), and NfL (ΔR^2^ = 0.018–0.022, p < 0.05).

Variance shared between age and white-matter integrity (age ∩ SLF) was significant across all six models (ΔR^2^ = 0.022–0.029, p_boot_ < 0.03), as was variance shared between age and each of the three plasma biomarkers (age ∩ BM; ΔR^2^ = 0.017–0.030, p_boot_ < 0.03), as expected given the mediation effects reported above. By contrast, variance shared between white-matter integrity and plasma biomarkers (SLF ∩ biomarker) was negligible and non-significant in all models (ΔR^2^ = −0.007 to 0.000, all p_boot_ > 0.12), providing direct evidence that the structural and molecular pathways contribute independently to sustained attention. Triple-shared variance was also non-significant across all models (all p_boot_ > 0.05).

To further disentangle AD-specific from nonspecific neuroaxonal contributions, we fit “full” models entering age, SLF integrity, NfL, and either pTau-181 or pTau-217 simultaneously (**Fig. S5**). In the pTau-181 models, both SLF FA and MD models explained significant variance in Att-Z (SLF FA: R^2^ = 0.204, F(7,138) = 5.06, p < 0.001; SLF MD: R^2^ = 0.178, F(7,138) = 4.26, p < 0.001), and age (β = −0.177 to −0.183, p < 0.05), SLF microstructure (SLF FA: β = +0.244, p < 0.01; SLF MD: β = −0.174, p < 0.02), and pTau-181 (β = −0.187 to −0.208, p < 0.02) remained independent predictors. NfL was no longer a significant independent predictor (β = −0.070 to −0.092, p = 0.156–0.223) after accounting for shared variance with pTau-181. Bootstrapped commonality analysis confirmed that pTau-181 and NfL shared significant overlapping variance in Att-Z (SLF FA model: ΔR^2^ = −0.023, p < 0.05; SLF MD model: ΔR^2^ = −0.022, p < 0.05), whereas unique SLF microstructure retained a significant independent contribution in both models (SLF FA: ΔR^2^ = −0.018, p < 0.01; SLF MD: ΔR^2^ = −0.019, p < 0.01; **Fig. S6**).

In the pTau-217 models, both SLF FA and MD models again explained significant variance in Att-Z (SLF FA: R^2^ = 0.191, F(7,138) = 4.65, p < 0.001; SLF MD: R^2^ = 0.161, F(7,138) = 3.79, p < 0.001). Age remained significant in both models (β = −0.182 to −0.192, p < 0.02), as did SLF FA (β = +0.240, p < 0.01) and SLF MD (β = −0.157, p < 0.05). Neither pTau-217 (β = −0.123 to −0.135, p = 0.060–0.076) nor NfL (β = −0.095 to −0.114, p = 0.107–0.154) retained significance when entered together, again suggesting that pTau-217 and NfL explain overlapping variance in attention that is more difficult to disentangle in a joint model in our CU sample and at this sample size. Commonality analysis corroborated this: shared pTau-217 ∩ NfL variance was significant in both models (SLF FA: ΔR^2^ = −0.023, p < 0.05; SLF MD: ΔR^2^ = −0.021, p < 0.05), while unique SLF microstructure again showed a significant independent contribution (SLF FA: ΔR^2^ = −0.014, p < 0.05; SLF MD: ΔR^2^ = −0.014, p < 0.05; **Fig. S7**).

Taken together, the commonality analyses converge with the mediation findings to support dissociable white-matter microstructural and AD-specific or neurodegeneration-related molecular contributions to age-related variance in sustained attention.

## Discussion

Differences among older adults in sustained attention are not reducible to a single neurobiological process. Here, combining DTI tractography of the SLF with plasma biomarkers measuring AD-related pathology and neurodegeneration, we show that SLF microstructural integrity and plasma pTau isoforms account for individual differences in attention and independently mediate age-related variance in sustained attention among CU older adults. These pathways contribute negligible shared variance to attentional performance, consistent with mechanistically independent routes through which these factors impact sustained attention.

The present findings implicate microstructural deterioration of the SLF as a structural mechanism linking ageing to impaired sustained attention. Lower FA and higher MD provide convergent indices of underlying white matter deterioration that together compromise the fidelity and speed of long-range prefrontal-parietal signal transmission. The consequence is precisely what gradCPT captures: attentional set degrades more rapidly over a sustained block, producing more frequent attentional lapses (higher RTV) and reduced capacity to discriminate goal-directed targets from lures (lower d’). Critically, when entering FA or MD from the SLF, CST, and CGC simultaneously, only SLF uniquely predicted all three attention indices, with bootstrap-confirmed coefficients significantly exceeding those of both control tracts. Age-to-attention mediation models further revealed that SLF partially mediates the effects of age on sustained attention. The frontoparietal regions connected by the SLF constitute core nodes of both the DAN, which supports voluntary top-down attention and the sustained maintenance of attentional engagement^[25]^, and the cognitive control network (CCN), which supports flexible goal maintenance and distractor suppression^[72]^; whether the effects observed here reflect disrupted communication within the DAN, the CCN, or both remains an open question that future work combining tractography with functional connectivity measures could address.

Plasma pTau-181 and pTau-217 also each mediated effects of age on sustained attention, independent of SLF integrity; plasma pTau was not associated with SLF FA or MD, and shared variance between SLF metrics and pTau was negligible across all analyses. These data suggest that AD-related effects on sustained attention are not mediated by alterations in SLF microstructure, and likely impact attention through other pathways not measured here. Given plasma pTau measures, and in particular pTau-217, have been linked to abnormal amyloid accumulation^[39,42,45]^, one possibility is that these effects reflect the impact of amyloid plaques in frontal and parietal association cortices^[23,24]^ on the cortical machinery of top-down attentional control. Alternatively, the observed effects may be linked to early regional tau accumulation among CU, given that individuals with elevated plasma pTau biomarkers are also more likely to exhibit elevated tau PET signal^[41,47]^. The locus coeruleus (LC), a brainstem nucleus that is the brain’s principal source of norepinephrine^[21]^, is among the earliest structures to accumulate pre-tangle tau pathology in the AD cascade, predating neocortical tau spread by decades^[20,50]^. Notably, the LC provides diffuse noradrenergic innervation to prefrontal and parietal cortices, supporting sustained attention through tonic arousal and phasic target detection^[73]^. Progressive LC neuronal loss would therefore be expected to produce precisely the gradCPT profile observed here: more frequent attentional lapses resulting in variable responding and reduced target discriminability. Although we did not measure LC integrity directly, circulating pTau-181 and pTau-217 show convergent associations with multiple indices of LC pathology, including LC neuron count at autopsy^[74]^ and LC neuromelanin MRI signal in living adults^[75]^. Future studies measuring LC integrity via neuromelanin-sensitive MRI alongside plasma AD biomarkers or more direct PET measures of tau accumulation and sustained attention would provide a critical test of this proposed pathway.

Interestingly, plasma NfL was also negatively associated with sustained attention performance and mediated effects of age on performance, independent of effects of SLF integrity. However, this effect did not remain significant when modelled jointly with pTau-181 or pTau-217, suggesting shared variance between these factors. Given that neuronal injury is a downstream effect of abnormal AD pathology^[17,53]^, this pattern is consistent with the possibility that these biomarkers measure different aspects of a common pathway, rather than exerting independent effects. However, due to the modest sample size and restricted range in these plasma markers among CU older adults, we interpret the effects of these combined models with caution. Additional work is needed to determine potential effects of neuronal injury, which can arise due to both AD-related and AD-independent processes, on attention. In contrast, GFAP, an index of astrocytic reactivity that rose significantly with age, showed no association with sustained attention in any model. This finding suggests that astrocyte reactivity does not explain variance in sustained attention, at least among CU older adults. Future research leveraging additional markers in populations with greater AD burden may provide complementary insights into the potential contributions of astrocyte reactivity and/or other alterations in innate immune processes to sustained attention impairment.

The mechanistic independence of the two pathways carries direct translational implications. The SLF pathway is substantially shaped by vascular health, myelination, sleep architecture, and aerobic fitness^[76,77]^, all potentially modifiable before clinical impairment and targetable independently of AD pathology. The molecular pathway is anchored in AD-related pathological accumulation, suggesting that strategies augmenting noradrenergic tone or tau-directed therapeutics may act preferentially on this route. CU individuals with degraded SLF but low plasma pTau or with elevated pTau but preserved SLF integrity may represent different biological profiles of attentional vulnerability. This argues for integrating white-matter microstructure alongside AD biomarkers in cognitive ageing frameworks, particularly in CU populations where the dissociation between structural and molecular axes is likely most pronounced.

Several limitations warrant acknowledgment. The cross-sectional design precludes causal inference about temporal ordering and does not permit examination of age-related decline per se; rather, observed age associations reflect cross-sectional variance across CU individuals differing in age. Longitudinal data are required to establish directionality and determine whether the two pathways diverge or converge over the lifespan. Tractography was limited to measuring FA and MD; advanced microstructural models such as NODDI^[78]^ could disambiguate myelin- and axon-density contributions to SLF integrity. The plasma subsample (N = 146) is modest for parallel mediation; replication in larger independent cohorts is warranted. Although plasma pTau-181 and pTau-217 are robust markers of underlying AD pathology, they are correlated with but do not provide direct measures of regional amyloid or tau accumulation; PET imaging would offer greater insight into the regionally specific effects of AD pathology on the cortical networks supporting attention. Finally, restriction to CU older adults limits biomarker and cognitive range; effect sizes are expected to be larger across the clinical continuum. This restriction is, however, also one of the study’s clearest strengths: demonstrating dissociable structural and molecular pathways before clinical impairment identifies them as targets precisely at the stage when intervention remains most tractable.

## Conclusion

Individual differences and age-related variability in sustained attention in cognitively unimpaired older adults reflect two mechanistically distinct and dissociable neurobiological pathways: microstructural deterioration of the frontoparietal SLF, which degrades the structural connections supporting prefrontal-parietal attentional signalling, and accumulation of AD-related pathology indexed by plasma pTau, which may erode sustained attention through disruption of neuromodulatory systems and progressive involvement of the cortical substrates supporting top-down attentional control. By contrast, effects of neuroaxonal injury (NfL) on attention were attenuated when modelled alongside pTau, consistent with neuroaxonal injury as a downstream consequence of AD pathology rather than an independent driver. Whether NfL reflects a mediating step within the AD pathway warrants examination in larger samples with greater biomarker range. The observed mechanistic independence of SLF microstructure and AD-related pathology identifies two separable biological targets for preserving attentional capacity before clinical impairment emerges.

## Acknowledgements

We thank the members of the Wagner and Mormino laboratories for discussions and support. We are grateful to I. Sai and A. Romero for assistance with data collection, and to the AMASS and SAMS volunteers for participation in the study.

## Author contributions

Conceptualization: S.S. and A.D.W. Methodology: S.S., J.P., J.R.B., A.N.T., A.D.W., and E.C.M. Software: S.S. Validation: S.S., A.D.W., A.N.T., and E.C.M. Formal analysis: S.S. Investigation: S.S., J.P. & J.R.B. Resources: A.D.W. and E.C.M. Data curation: S.S., J.P and J.R.B. Visualization: S.S. Supervision: A.D.W. and E.C.M. Project administration: A.D.W., E.C.M., and A.N.T. Funding acquisition: A.D.W. and E.C.M. Writing-original draft: S.S. Writing-review and editing: A.D.W., A.N.T., E.C.M., S.S., and J.S. All authors reviewed the manuscript. Authors J.P. and J.R.B contributed equally to the manuscript and are listed alphabetically.

## Data and materials availability

All data needed to evaluate the conclusions in this paper are present in the paper and/or the Supplementary Materials.

