## Supplementary Material for "Dissociable structural and molecular pathways of age-related variability in sustained attention"

### Supplementary Section

#### S1: Hemispheric SLF FA and MD Associations with Sustained Attention

To examine whether associations between SLF microstructural integrity and sustained attention were lateralised, right and left hemisphere SLF FA and MD were examined separately (**Figs. S1 & S2**). Both hemispheres independently predicted sustained attention performance across both microstructural metrics. Left SLF FA positively predicted d′ ($\beta = +0.230$, $p < 0.01$), negatively predicted RTV ($\beta = -0.205$, $p < 0.01$), and positively predicted Att-Z ($\beta = +0.233$, $p < 0.01$); left SLF MD showed a consistent pattern: d′ ($\beta = -0.189$, $p < 0.01$), RTV ($\beta = +0.203$, $p < 0.01$), and Att-Z ($\beta = -0.210$, $p < 0.01$). Similarly, right SLF FA significantly predicted d′ ($\beta = +0.238$, $p < 0.001$), RTV ($\beta = -0.243$, $p < 0.001$), and Att-Z ($\beta = +0.257$, $p < 0.001$); and right SLF MD showed a consistent pattern: d′ ($\beta = -0.175$, $p < 0.05$), RTV ($\beta = +0.132$, $p < 0.05$), and Att-Z ($\beta = -0.164$, $p < 0.05$). Bootstrap comparisons confirmed no significant hemispheric differences between left and right FA coefficients nor between left and right MD coefficients for any attention outcome (all $p > 0.35$), supporting a bilateral SLF contribution to sustained attention without significant hemispheric asymmetry.

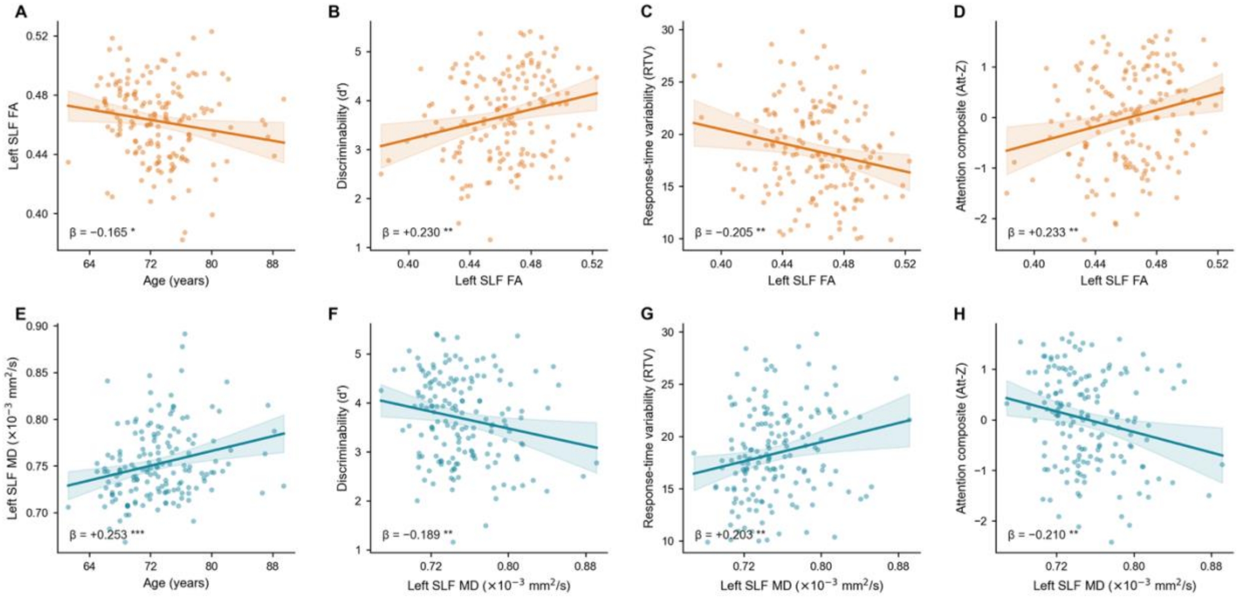


***Figure S1.*** *Partial-regression associations (N = 162) between left hemisphere SLF fractional anisotropy (FA; orange, top row) or mean diffusivity (MD; teal, bottom row) and age (A, E) or sustained attention (B–D, F–H). Older age was associated with lower left SLF FA and higher left SLF MD. Higher left SLF FA and lower MD were associated with better gradCPT discriminability (d′; B, F), lower response-time variability (RTV; C, G), and a higher attention composite score (Att-Z; D, H). Each panel plots observed outcome values against the Age or DTI predictor after residualising both variables on covariates. Lines and shading show OLS predicted values ± 95% CI. Standardised β coefficients and significance symbols refer to the partial regression of the Age or DTI metric on the outcome. *p < 0.05, **p < 0.01, ***p <0.001 (one-tailed).*

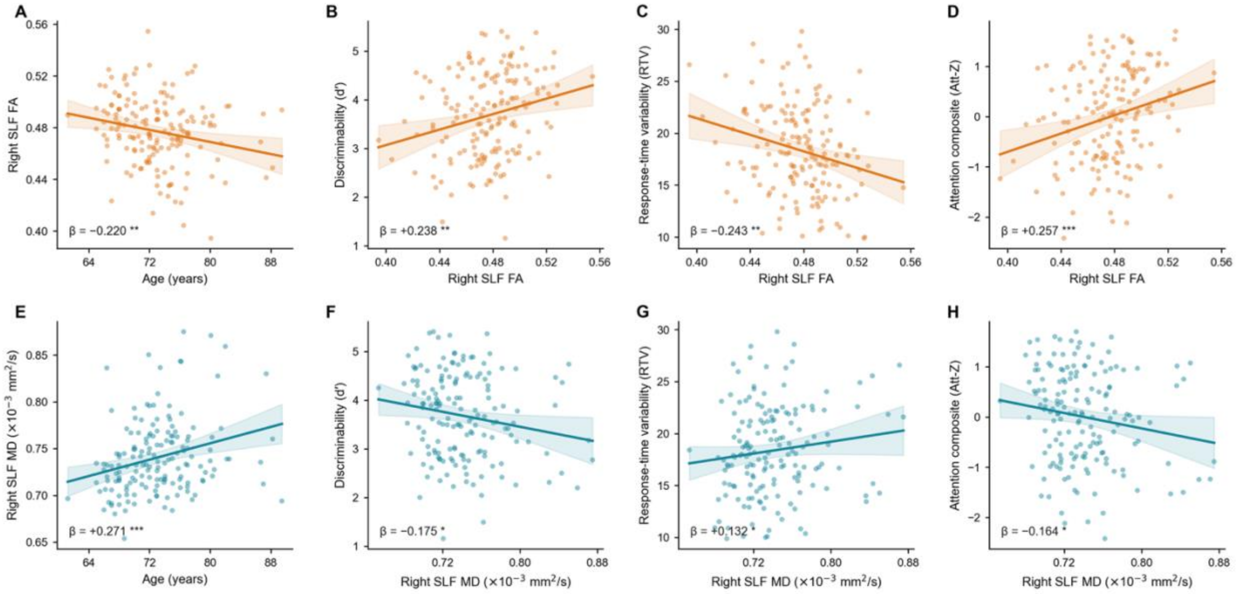


***Figure S2.*** *Partial-regression associations (N = 162) between right hemisphere SLF fractional anisotropy (FA; orange, top row) or mean diffusivity (MD; teal, bottom row) and age (A, E) or sustained attention (B–D, F–H). Older age was associated with lower right SLF FA and higher right SLF MD. Higher right SLF FA and lower MD were associated with better gradCPT discriminability (d'; B, F), lower response-time variability (RTV; C, G), and a higher attention composite score (Att-Z; D, H). Each panel plots observed outcome values against the Age or DTI predictor after residualising both variables on covariates. Lines and shading show OLS predicted values ± 95% CI. Standardised β coefficients and significance symbols refer to the partial regression of the Age or DTI metric on the outcome. *p < 0.05, **p < 0.01, ***p < 0.001 (one-tailed).*

### S2: Plasma Biomarkers and SLF Microstructural Integrity

To test whether plasma biomarker burden predicts SLF white-matter microstructure, and thereby whether plasma pathology might act on sustained attention via structural degradation of frontoparietal pathways, we regressed bilateral SLF FA and MD on each of the four primary plasma biomarkers, controlling for age, sex, education, mean framewise displacement. (**Fig. S3**).

None of the four biomarkers significantly predicted bilateral SLF FA: pTau-181 ($\beta = +0.023$, $p = 0.396$), pTau-217 ($\beta = -0.005$, $p = 0.475$), NfL ($\beta = +0.073$, $p = 0.202$), and GFAP ($\beta = -0.110$, $p = 0.097$; all $p_{FDR} > 0.1$). For SLF MD, results were similarly null for pTau-181 ($\beta = -0.106$, $p = 0.109$), pTau-217 ($\beta = -0.030$, $p = 0.359$), and NfL ($\beta = -0.024$, $p = 0.393$); GFAP showed a marginal positive association ($\beta = +0.121$, $p = 0.074$) that did not survive FDR correction.

To formally test whether plasma biomarkers mediate age-related SLF microstructural decline, we estimated bootstrap mediation models (5,000 samples) with age as the predictor and bilateral SLF FA or MD as the outcome, controlling for education, sex, and mean framewise displacement (**Fig. S4**). No biomarker carried a significant indirect effect on SLF FA: pTau-181 ($p = 0.379$), pTau-217 ($p = 0.479$), NfL ($p = 0.162$), and GFAP ($p = 0.066$). Parallel models for SLF MD likewise yielded no significant indirect effects: pTau-181 ($p = 0.101$), pTau-217 ($p = 0.354$), NfL ($p = 0.377$), and GFAP ($p = 0.061$). Together, these findings indicate that pTau-181, pTau-217, and NfL plasma biomarker burden does not account for age-related SLF microstructural decline, consistent with the interpretation that structural and these molecular pathways to sustained attention operate independently. Given the trend-level relationships observed with GFAP, we do not draw a strong conclusion regarding the possible mediating role of GFAP. Future studies with even larger sample sizes are warranted to address this possibility.

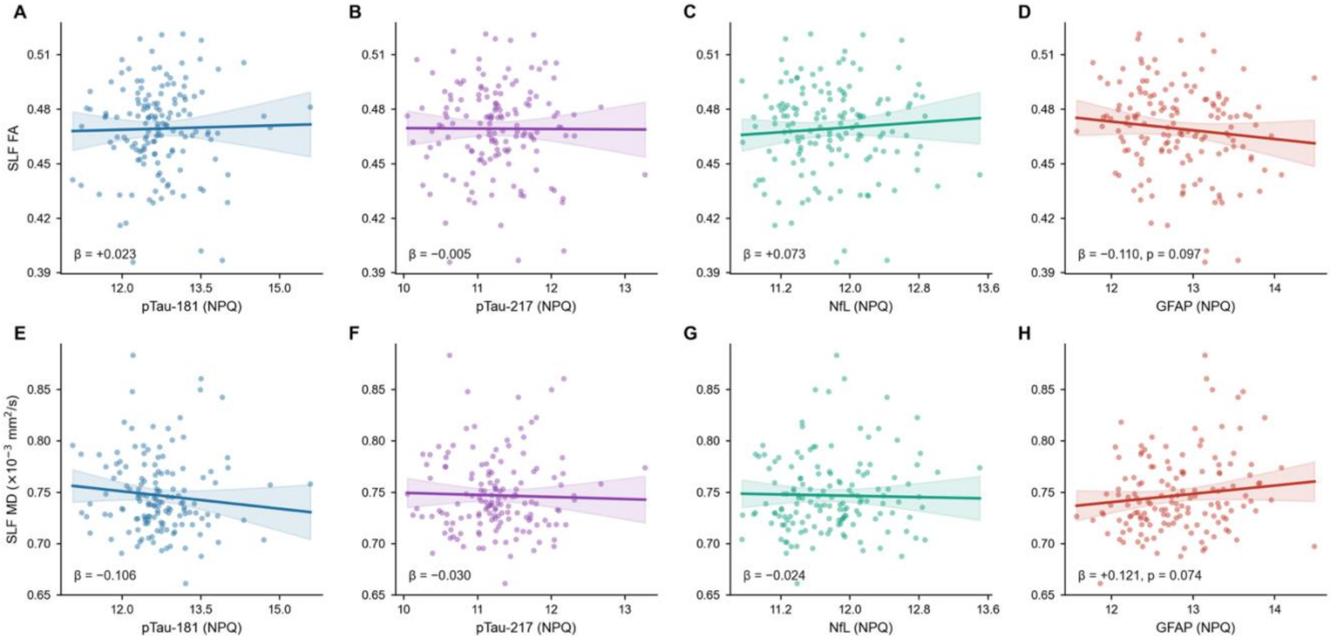


***Figure S3.*** *Associations between primary plasma biomarkers and bilateral SLF white-matter microstructure (N = 146). Top row (A–D): partial-regression associations between each plasma biomarker and SLF fractional anisotropy (FA). Bottom row (E–H): partial-regression associations between each plasma biomarker and SLF mean diffusivity (MD). Columns show pTau-181 (A, E), pTau-217 (B, F), NfL (C, G), and GFAP (D, H). Each panel plots observed outcome values against the biomarker predictor after residualising both variables on age, sex, education, and mean framewise displacement. Lines and shading indicate OLS model-predicted values ± 95% CI. Standardised β coefficients reflect the partial association between each biomarker and SLF microstructure, controlling for the same covariates. No association survived FDR correction; uncorrected p-values are shown where p < 0.10.*

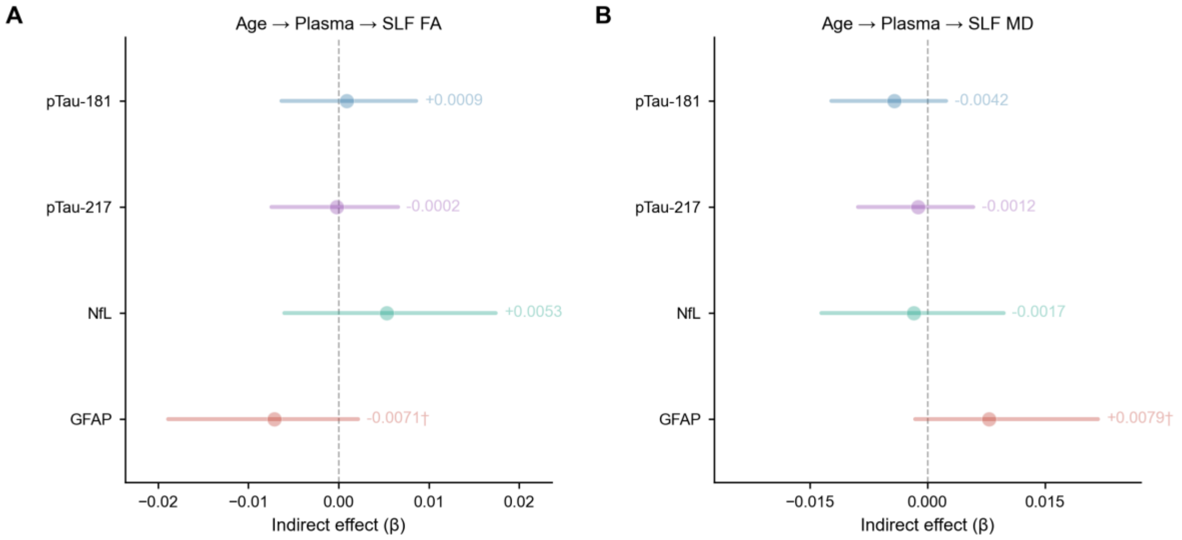


***Figure S4.*** *Bootstrap mediation: Age → plasma biomarker → SLF microstructural integrity (N = 146). Forest plots show indirect effects (a × b path product) with bias-corrected 95% bootstrap confidence intervals (5,000*

*samples) for mediation models in which each plasma biomarker was tested as a mediator of age-related change in bilateral SLF FA (A) or SLF MD (B). Prior to mediation analysis, SLF FA and MD were z-scored to place indirect effects on a common standardised scale and facilitate comparison across metrics. Covariates: sex, education, and mean framewise displacement. No biomarker carried a significant indirect effect on SLF FA or MD. GFAP showed trending indirect effects on both SLF FA and SLF MD, that did not reach significance. Faded markers indicate non-significant indirect effects; circles indicate non-significant, diamonds indicate significant ($p < 0.05$). †$p < 0.10$ (one-tailed).*

### S3: Full Commonality Analyses

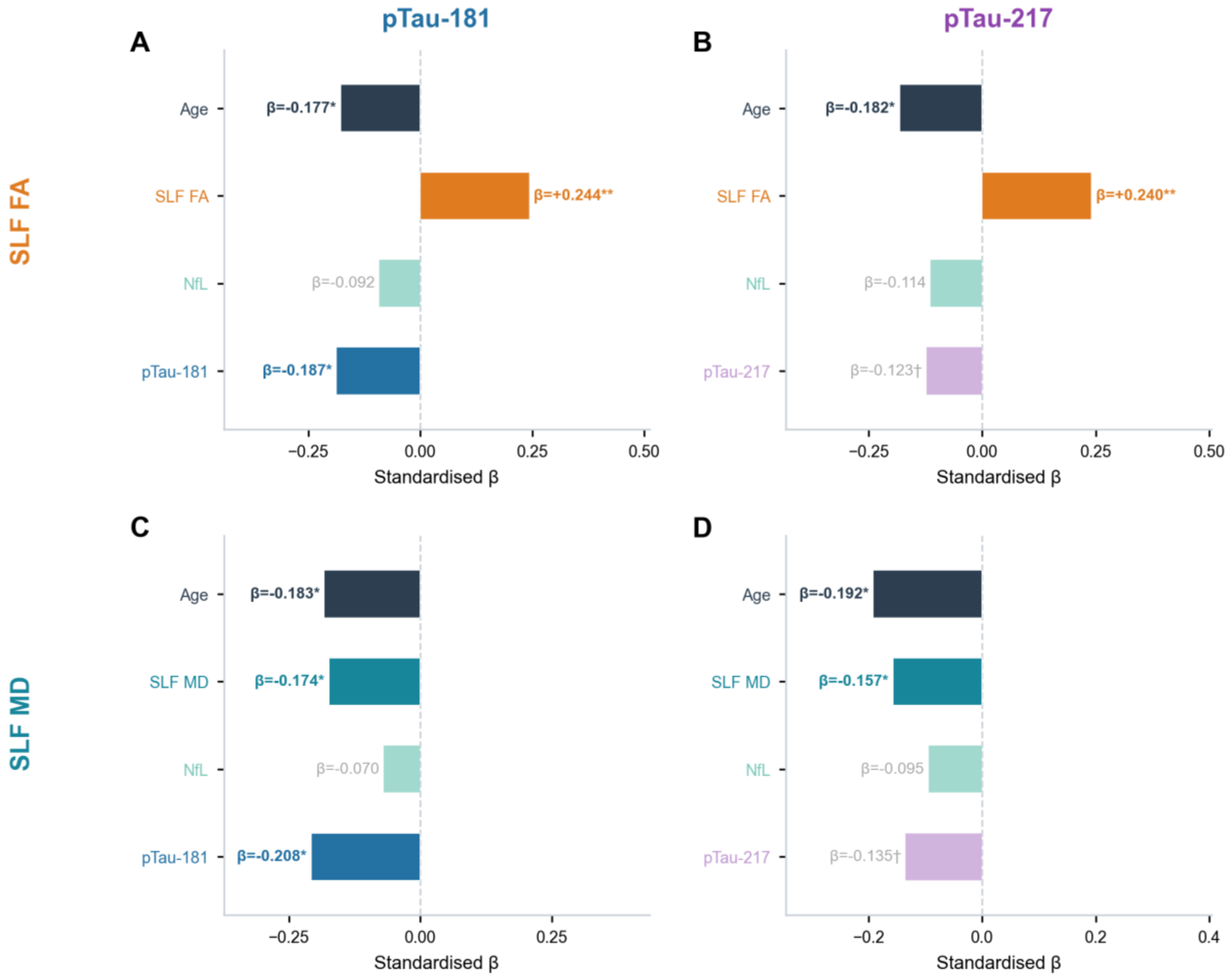


***Figure S5.*** *Full OLS models: Age, SLF microstructure, NfL, and pTau predictors of attention (N = 146). Standardised β coefficients from four OLS regression models predicting the Att-Z composite (higher = better attention), each including age, bilateral SLF FA (A, B) or SLF MD (C, D), NfL, and either pTau-181 (A, C) or pTau-217 (B, D) as predictors. Covariates: education, sex, and mean framewise displacement. P-values are one-tailed. Bars are fully opaque for $p < 0.05$; faded) for $p \geq 0.05$. *$p < 0.05$; **$p < 0.01$. †$p < 0.10$.*

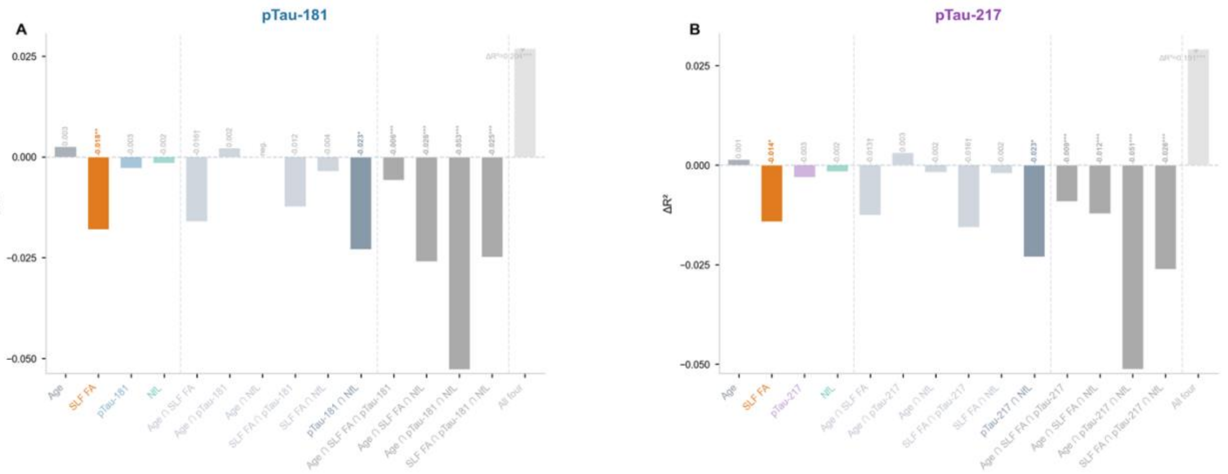


***Figure S6.*** *Full commonality decomposition: age + SLF FA + NfL + pTau → Att-Z (N = 146). Bars show all 15 bootstrapped ΔR² components (5,000 resamples, one-tailed p) for 4-predictor models with SLF FA as the white-matter metric and either pTau-181 (A) or pTau-217 (B). Covariates: education, sex, mean framewise displacement. Components are grouped (left to right) by unique predictors, pairwise shared, triple shared, and total (All four; annotated separately above axis). Opaque bars: $p < 0.05$; faded: $p \geq 0.05$. Unique SLF FA was the only predictor with a significant independent contribution in both models (pTau-181: $\Delta R^2 = -0.018$, $p < 0.01$; pTau-217: $\Delta R^2 = -0.014$, $p < 0.05$). Shared pTau ∩ NfL variance was significant in both models (pTau-181: $\Delta R^2 = -0.023$, $p < 0.05$; pTau-217: $\Delta R^2 = -0.023$, $p < 0.05$), indicating overlapping biomarker contributions that cannot be independently resolved at this sample size.*

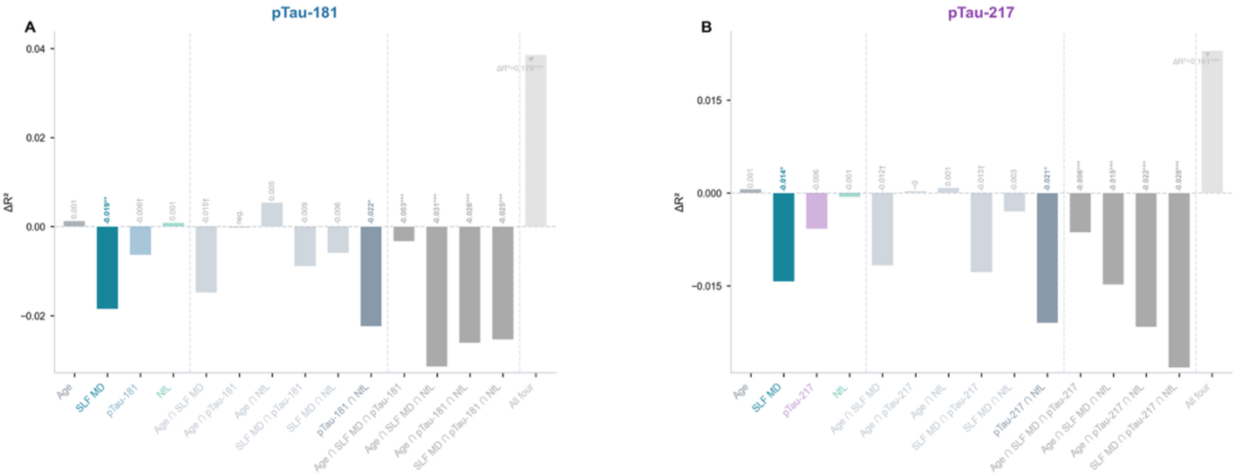


***Figure S7.*** *Same as Fig S6 but with SLF MD as the white-matter metric. Unique SLF MD was significant in both models (pTau-181: $\Delta R^2 = -0.019$, $p < 0.01$; pTau-217: $\Delta R^2 = -0.014$, $p < 0.05$). Shared pTau ∩ NfL variance was again significant (pTau-181: $\Delta R^2 = -0.022$, $p < 0.05$; pTau-217: $\Delta R^2 = -0.021$, $p < 0.05$).*
